# ScreenDMT reveals metabolites replicably associated with sleep disorders across three human cohorts

**DOI:** 10.64898/2026.10.10.755811

**Authors:** Jonathan M Dreyfuss, Dhruv Dubey, Yu Zhang, Michael R Brown, Jerome I Rotter, Stephen S Rich, Peter Y Liu, Susan Redline, Heming Wang, Robert E Gerszten, Clary B Clish, Annah B Wyss, Alanna C Morrison, Jan Bressler, Chloé Sarnowski, Bing Yu, Pamela L Lutsey, Brendan P Lucey, Jee-Young Moon, Hui Pan, Vera Djordjilovic, Tamar Sofer

## Abstract

Replication of scientific findings across independent studies is essential, yet it remains challenging in high-dimensional biological datasets. A critical aspect of true replication requires detecting not only the repeated presence but also the consistent direction of an effect. Traditional methods for evaluating replication often ignore effect directionality or lack adequate statistical power when adjusting for multiple comparisons. To address this, we extend the directional MaxP test (DMT) and our screening-integrated procedure, ScreenDMT, to evaluate directional replication across an arbitrary number of independent studies. We mathematically prove ScreenDMT’s validity for an arbitrary number of studies when the studies’ sample sizes grow unbounded. Through extensive simulation studies, we demonstrate that ScreenDMT effectively controls error rates, such as the false discovery rate (FDR), in finite sample sizes and yields higher statistical power than competing methods especially for three studies. We applied the extended ScreenDMT to untargeted blood metabolomics profiles across three diverse human cohorts—ARIC (N=1,337), MESA (N=1,874), and HCHS/SOL (N=6,180)—to discover replicable biomarkers for dichotomous insomnia, long sleep, short sleep, and excessive daytime sleepiness (EDS). We found low levels of cortisol to be replicably associated with EDS, and high levels of n-formylmethionine to be associated with long sleep. Furthermore, directional pathway analysis revealed that low levels of xanthine metabolites are replicably associated with insomnia. These convergent findings validate ScreenDMT as a powerful methodology for simultaneous multi-cohort replication and offer insights into the metabolic signatures underlying sleep disorders across diverse populations.

## Introduction

Replication across independent studies shows whether a new discovery is reliable and generalizable. When a hypothesis or many hypotheses are tested across multiple studies, they are typically analyzed with a meta-analysis, such as inverse-variance weighted meta-analysis, but such a meta-analysis can be significant due to a strong effect in only one study, so it does not ensure replication. One traditional replication approach is to designate a single discovery cohort (adjusting for multiple testing across all analytes) and evaluate top hits in a replication cohort with reduced multiple testing burden. However, results are sensitive to the choice of discovery cohort—which can be arbitrary—and this sequential framework does not scale naturally to multiple independent cohorts. Instead, we evaluate all cohorts simultaneously.

Rigorously assessing replication, particularly in high-dimensional biological datasets where numerous analytes are measured, presents substantial statistical challenges. A critical aspect of true replication is not only the repeated presence of an effect but also its concordant direction across different cohorts or physiologically-related conditions. Traditional methods for evaluating replication have often ignored this directionality, lost statistical power, or failed to control error rates when adjusting for multiple comparisons (Bogomolov and Heller 2023).

For testing a single feature or analyte for association with a phenotype across two independent studies, consider the example of testing the association between each subject’s plasma abundance of a lipid and that subject’s body mass index (BMI) across n=2 independent cohorts. Let the true but unknown values of the association parameters representing linear regression model coefficients in the two cohorts be and, and the null hypothesis per cohort be 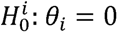. Across both cohorts, the null hypothesis representing no replication without regard to direction is that θ_1 = 0 or_ θ_2 = 0_ against the alternative that both θ_1 ≠ 0_ and θ_2 ≠ 0_. On a graph, the null would be true if and only if the parameter value sat on the x-axis or the y-axis (or both).

From each cohort we obtain the association’s summary statistics such as z-score and p-value, e.g. (Z_1_, P_1_) and (Z_2_, P_2_). A common test for replication in such cases that does not require a consistent direction is the joint significance or MaxP test whose p-value = max{P_1_, P_2_}=P_max_. This was found to control its false positive rate, have more power than some other common methods, and be the likelihood ratio test (LRT) (MacKinnon, Lockwood et al. 2002, Barfield, Shen et al. 2017, Liu, Shen et al. 2022). Note that some of these studies examined this test for the related problem of causal mediation. However, this test identifies analytes with both concordant or discordant signs.

When we account for direction, the null hypothesis of directional replicability in Equation 1 is that one of the effects is absent or their signs are different, whereas the alternative is that both effects are present and in the same direction.

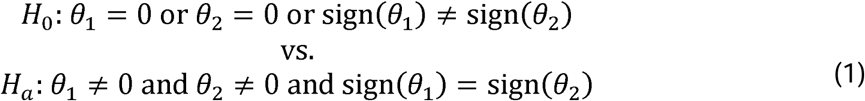

We previously introduced a directional MaxP test (DMT) that has more power for the directional replicability null hypothesis with two studies, and we showed that it is a LRT (Dreyfuss, Yuchi et al. 2021, Dreyfuss, Djordjilović et al. 2024). For replication, the DMT p-value is defined as in Equation 2.

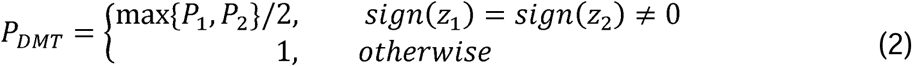

For testing multiple analytes for association with a phenotype across two independent studies, consider the example of testing the association of each lipid with BMI. We will have one DMT nominal p-value per lipid, and due to multiple testing, these DMT nominal p-values have to be adjusted. We could adjust with a standard multiple testing correction e.g. Benjamini and Hochberg’s false discovery rate (FDR) procedure. To improve power beyond standard adjustment, we integrated DMT with AdaFilter’s (Wang, Gui et al. 2022) screening procedure. AdaFilter defines a filtering and selection p-value per analyte, where the selection p-value cannot be less than the filtering p-value. For an analyte, consider *p_min =_* _min_{p_1,_ p_2_} to be the filtering p-value and *p_max =_* _max_{p_1,_ p_2_} to be the selection p-value. When not accounting for direction, the procedure filters out analytes whose filtering p-value is so large that their selection p-value cannot be statistically significant, and then only adjusts the selection p-values that survived filtering for multiple testing.

A naive integration would be to define the filtering p-value to be P_min_ and define the selection p-value to be the DMT p-value, but this is not valid, since the DMT p-value may be less than P_min_. Instead, we previously defined the selection p-value = max{P_max_/2, P_min_} for our ScreenDMT method, which we proved to be valid for two studies. We also found it to have more power than adjusting the DMT p-values without filtering in our simulations and application, where we identified that the lipid 9,10- diHOME’s abundance was replicated as being negatively associated with BMI, and we experimentally validated a potential mechanism (Dreyfuss, Djordjilović et al. 2024).

Here, we extend the testing procedures DMT and ScreenDMT to testing replication across more than two studies. We apply them to identify metabolites replicably associated with the sleep disorders insomnia, long sleep, short sleep, and excessive daytime sleepiness (EDS) across three human cohorts. Disordered sleep affects about a third of the U.S. population and is linked to many negative health and societal outcomes, but its biological mechanisms remain poorly understood (Institute of Medicine Committee on Sleep Medicine and Research 2006). To identify biomarkers that could elucidate etiology and provide therapeutic targets, a promising approach is blood metabolomics. We previously identified metabolites associated with EDS in the Hispanic Community Health Study/Study of Latinos (HCHS/SOL) and tested the significant metabolites for nominal significance in Multi-Ethnic Study of Atherosclerosis (MESA) and in two other cohorts that only had proxy metabolites, which yielded metabolites involved in steroid metabolism (Faquih, Potts et al. 2025). To our knowledge, this is the first simultaneous replication test of sleep disorder associated metabolites across three human cohorts.

## Results

### Extension of DMT to an arbitrary number of studies

When testing whether an analyte’s association with an outcome replicates across *n* out of *n* studies, for arbitrary *n*, the MaxP test p-value is P_max_ = max{P_1_, P_2_, …, P_n_}. We show in the supplement that this is the likelihood ratio test (LRT) for the non-directional null (Proposition, Note S1).

To test the directional null for replication across *n* out of *n* studies, we define the DMT p-value as:

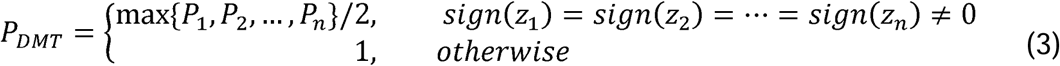

A proof that the DMT p-value is the LRT for the case when the signs are non-zero and the same has been given before (Sasabuchi 1980, Berger 1989).

Another approach to test the directional null that was adapted from a meta-analysis technique introduced by Pearson, which we term “Pearson”, involves calculating the MaxP for each analyte to right-sided p-values of all studies, then applying MaxP to left-sided p-values, and doubling the minimum from the two applications (Owen 2009, Wang, Gui et al. 2022). This Pearson-based procedure also underlies AdaFilter’s directional approach.

### Extension of ScreenDMT to an arbitrary number of studies

For an arbitrary number of studies *n* with the non-directional null, AdaFilter’s (Wang, Gui et al. 2022) screening procedure for an analyte is defined in terms of its ordered p-values, where the smallest p-value across studies is denoted P_(1)_, the next smallest p-value is P_(2)_, all the way up to P_(n)_. The selection p-value is P_(n)_, which is P_max_, and the filtering p-value is the next least significant p-value, P_(n-1)_. So for the directional null, our filtering p-value is similarly P_(n-1)_ but our selection p-value, which must not be less than P_(n-1)_, is max{P_(n-1)_, P_max_/2}. For the screening to be valid we must demonstrate a form of conditional validity per metabolite where the conditional distribution of the selection p-value given the filtering p-value is either uniform or stochastically dominates the uniform distribution. Formally, this is

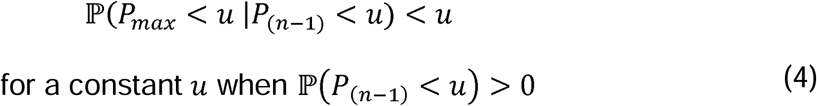

We have a closed form expression for this conditional probability, which we use to prove that the ScreenDMT procedure satisfies this conditional validity requirement asymptotically for an arbitrary number of studies *n,* i.e. in the limit where each study’s sample size increases without bound (Theorem, Note S1).

To see the validity when the studies have finite sample sizes, we examine our closed form expression in several scenarios (Figure 1). The conditional probability depends on the choice of *u* and on the association parameters standardized by the standard error, i.e. the delta parameters, which affect the distribution of the p-values. We plot this conditional probability case where there are three studies and the standardized association parameter for study one is δ_1_=0, δ_2_ is defined in the facet labels, and δ_3_is defined on the x-axis. We find that the conditional probability is slightly inflated when the delta parameters are positive and small, but when the delta parameters grow larger (as happens when the theta parameters are positive and the sample size grows) the conditional p-value is controlled, which is consistent with our theorem and what we saw previously for n=2 studies (Dreyfuss, Djordjilović et al. 2024).

**Figure 1:**
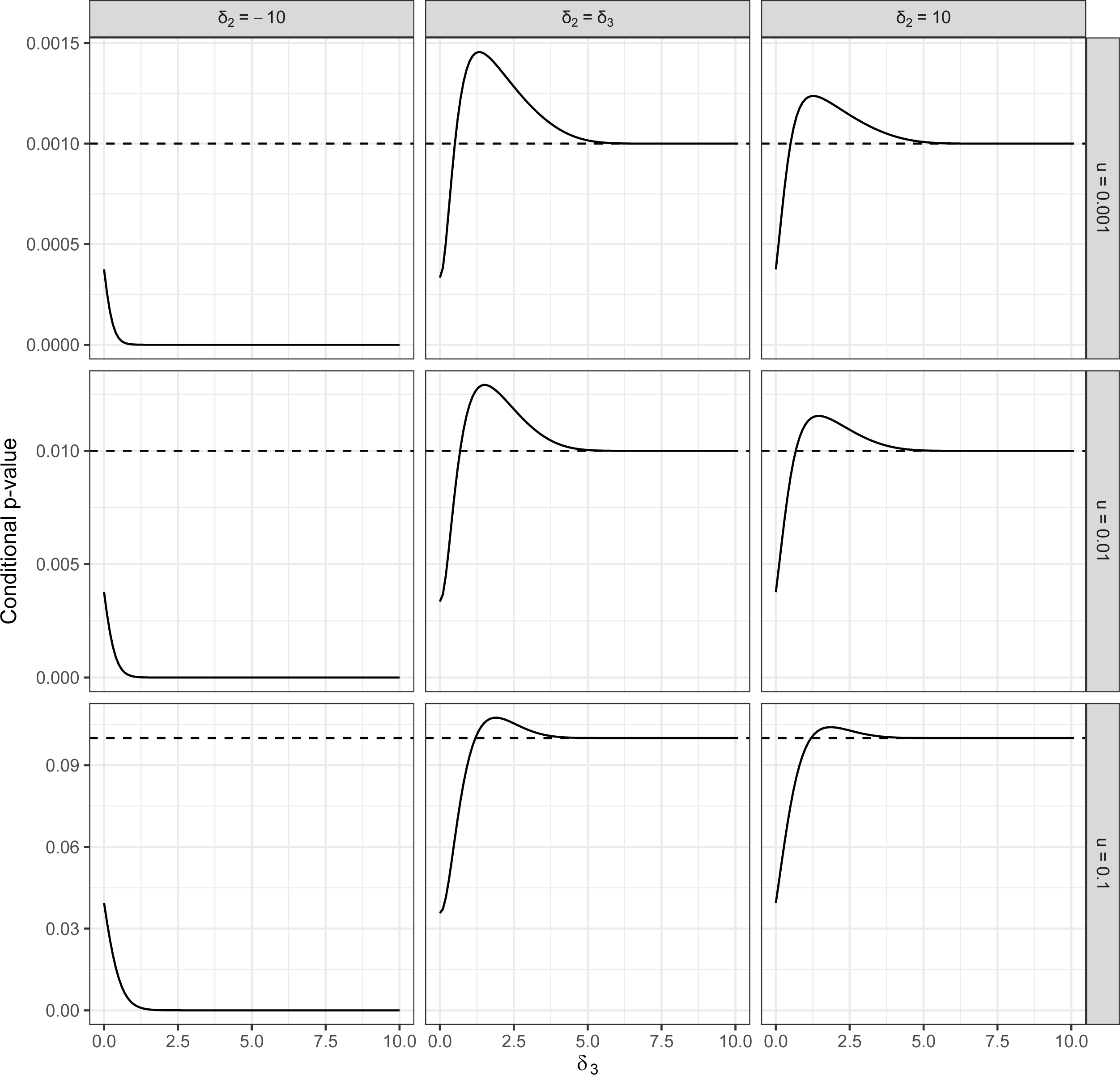
Behavior of conditional p-value in finite sample sizes. The y-axis represents the conditional p-value in 3 studies, which is P(p_DMT_ < u | p_(2)_ < u) < u where the true standardized effect sizes are δ_1_=0, δ_2_ as defined in the facet labels, and δ_3_as defined on the x-axis.

### Simulation Studies

Similarly to our previous work on ScreenDMT with n=2 studies (Dreyfuss, Djordjilović et al. 2024), we validate DMT and ScreenDMT for multiple studies via simulation across a variety of settings to demonstrate that they control the error rates and in some settings have more power than competing approaches. We conducted simulations with n=3 studies and separately with n=4 studies, where each study was simulated to contain measurements for the same 1,000 analytes, such as genes, proteins, or metabolites. Each element of the 1,000 x *n* matrix was simulated as signal (e.g. 3.5, −3.5) or no signal (zero) plus random noise as a random draw from the standard normal (z-score) distribution.

In each simulation, we simulated 5% of metabolites to be truly directionally replicated across all studies, meaning these metabolites had a nonzero signal of the same sign in every study (either all positive or all negative), whereas 95% of the metabolites were simulated under the directional null. We varied the proportion of metabolites under the directional null that were simulated with nonzero signal in at least one study across settings. There are many possible configurations of these signal profiles, as described in the Methods.

We evaluated both an equal-signal setting, where all nonzero signals have magnitude 3.5, and an unequal-signal setting, where nonzero signal magnitudes per metabolite varied across studies while keeping the overall average signal magnitude the same (if there’s an odd number of studies, in one study the signal magnitude is still 3.5). To evaluate robustness to dependence across metabolites, we simulated both weak and strong dependence using a block correlation structure, where metabolites within a block were correlated (correlation coefficient of 0.5) and block sizes were set to 10 or 100, respectively, like the AdaFilter simulations. Across all settings, we compared ScreenDMT and DMT to the competing methods AdaFilter (Wang, Gui et al. 2022), Pearson (Owen 2009), and Joint Mirror (Deng, He et al. 2024). Joint Mirror identifies which metabolites significantly replicate but does not calculate an FDR per metabolite. We evaluated power and error rate control at a 15% target for both FDR and family-wise error rate (FWER).

The simulation results for n=3 studies are shown in Figure 2. The x-axis represents the percent of analytes that do not truly directionally replicate across all n=3 studies. These analytes have different signal profiles across the 3 studies, e.g. one may have signal in only one or two studies, another may show signals of opposite direction across all 3 studies. Here, the signal is −3.5 or +3.5. These different signal profiles are sampled with equal probability. Five percent of the analytes are defined to truly directionally replicate across all 3 studies by having only positive or only negative effects. The remaining analytes do not have any nonzero signal in any study.

**Figure 2:**
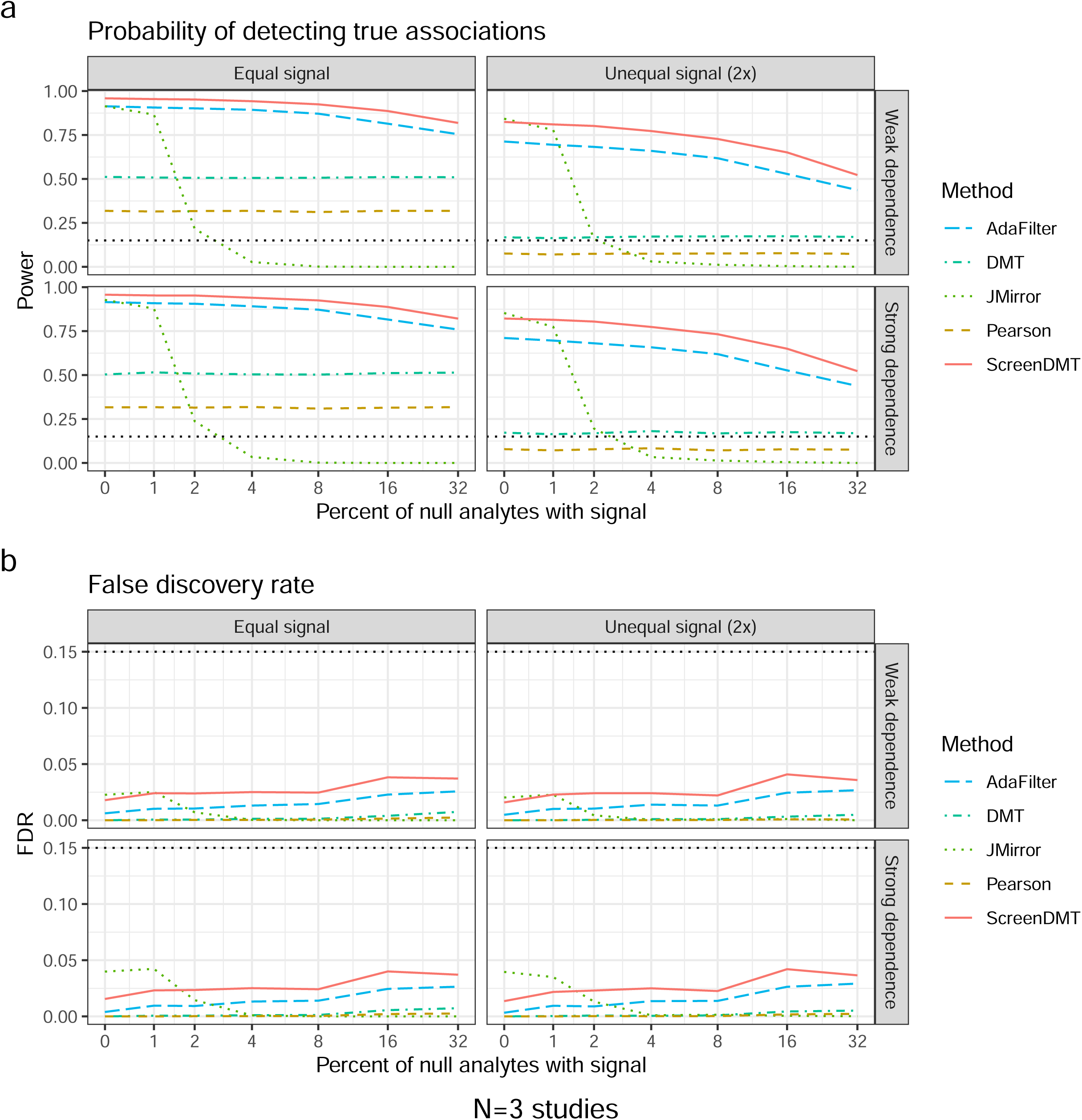
Power and FDR for multiple methods with 3 studies. The x-axis represents the percent of analytes that do not truly directionally replicate across all N studies but show effect in a subset of the studies or show effects of opposite direction in all N studies. Five percent of the analytes truly directionally replicate across all N studies by having only positive or only negative effects. The remaining analytes do not have any true effect in any study. (A) shows the power and (B) shows the false discovery rate (FDR).

Figure 2A shows the power, which is the estimated probability of detecting true associations out of the 1,000 simulations per setting, where we can see that DMT has more power than Pearson, and ScreenDMT has more power than any other method across nearly all settings. The only setting where ScreenDMT does not dominate is in the unequal signal setting where the x-value is exactly zero, which means that the only analytes with nonzero signal in any study happen to directionally replicate across all studies, since the Joint Mirror procedure has slightly more power. However, ScreenDMT shows much more power than the Joint Mirror procedure across most of the x-axis values and moderately more power than AdaFilter across the entire x-axis. In Figure 2B, we can see that all methods included here control their FDR at the nominal 15% level in all settings; in fact, they control their FDR at a much lower rate due to being over-conservative when testing the composite null.

In Figure 3A, we see simulation results for N=4 studies. ScreenDMT has more power than AdaFilter where there are fewer null analytes with nonzero signal, whereas it has more power than the Joint Mirror when there are not too few null analytes with nonzero signal. Overall, which of these three methods has the most power depends strongly on the proportion of null analytes with nonzero signal. In Figure 4, we show simulation results with FWER instead of FDR, where ScreenDMT is the most powerful when there are few null analytes with nonzero signal, whereas AdaFilter is most powerful otherwise, but the difference is small for an appreciable range of the x-axis. All methods control their error rates in Figures 3 and 4.

**Figure 3:**
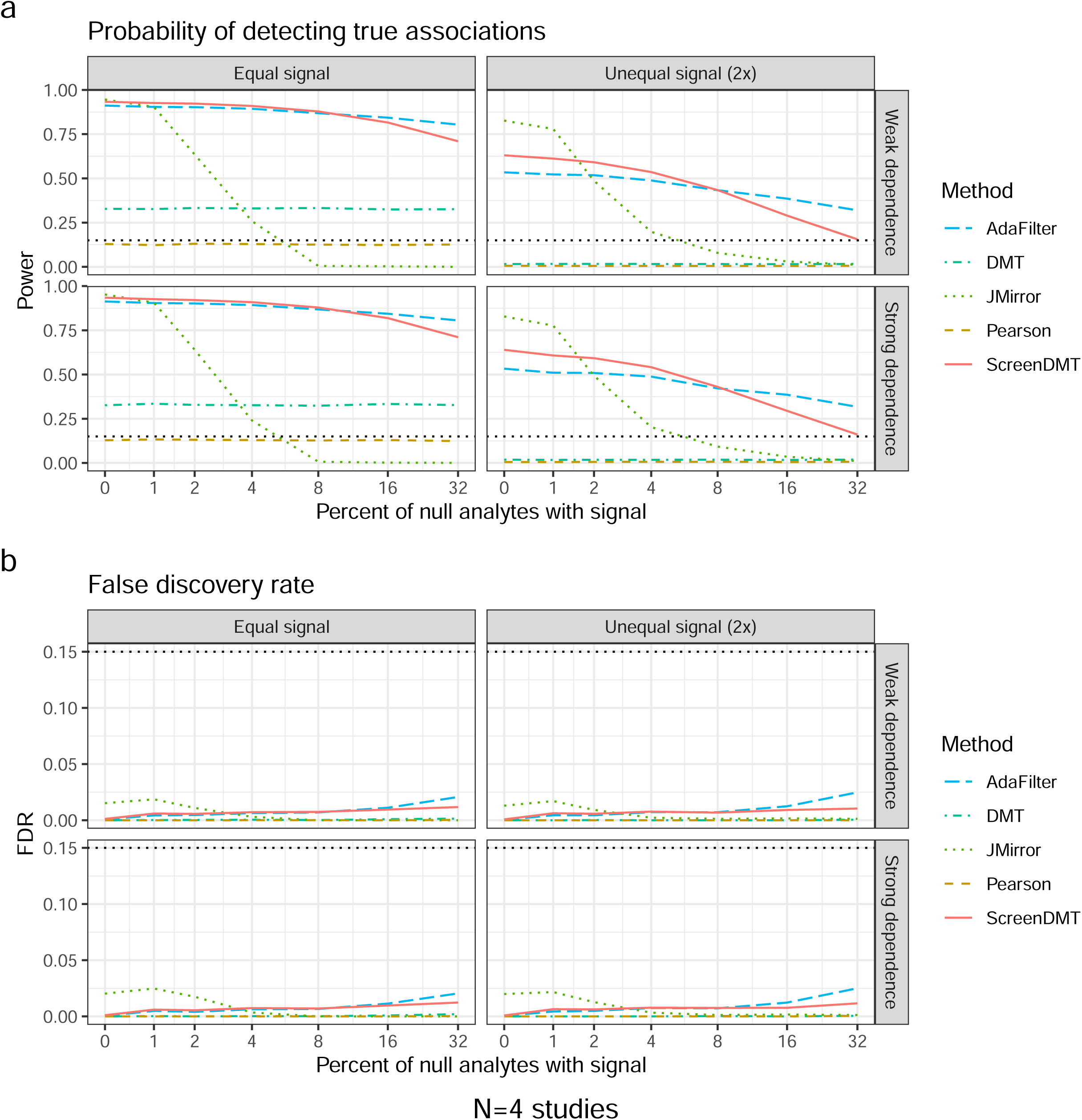
Power and FDR for multiple methods with 4 studies. The x-axis represents the percent of analytes that do not truly directionally replicate across all N studies but show effect in a subset of the studies or show effects of opposite direction in all N studies. Five percent of the analytes truly directionally replicate across all N studies by having only positive or only negative effects. The remaining analytes do not have any true effect in any study. (A) shows the power and (B) shows the FDR.

**Figure 4:**
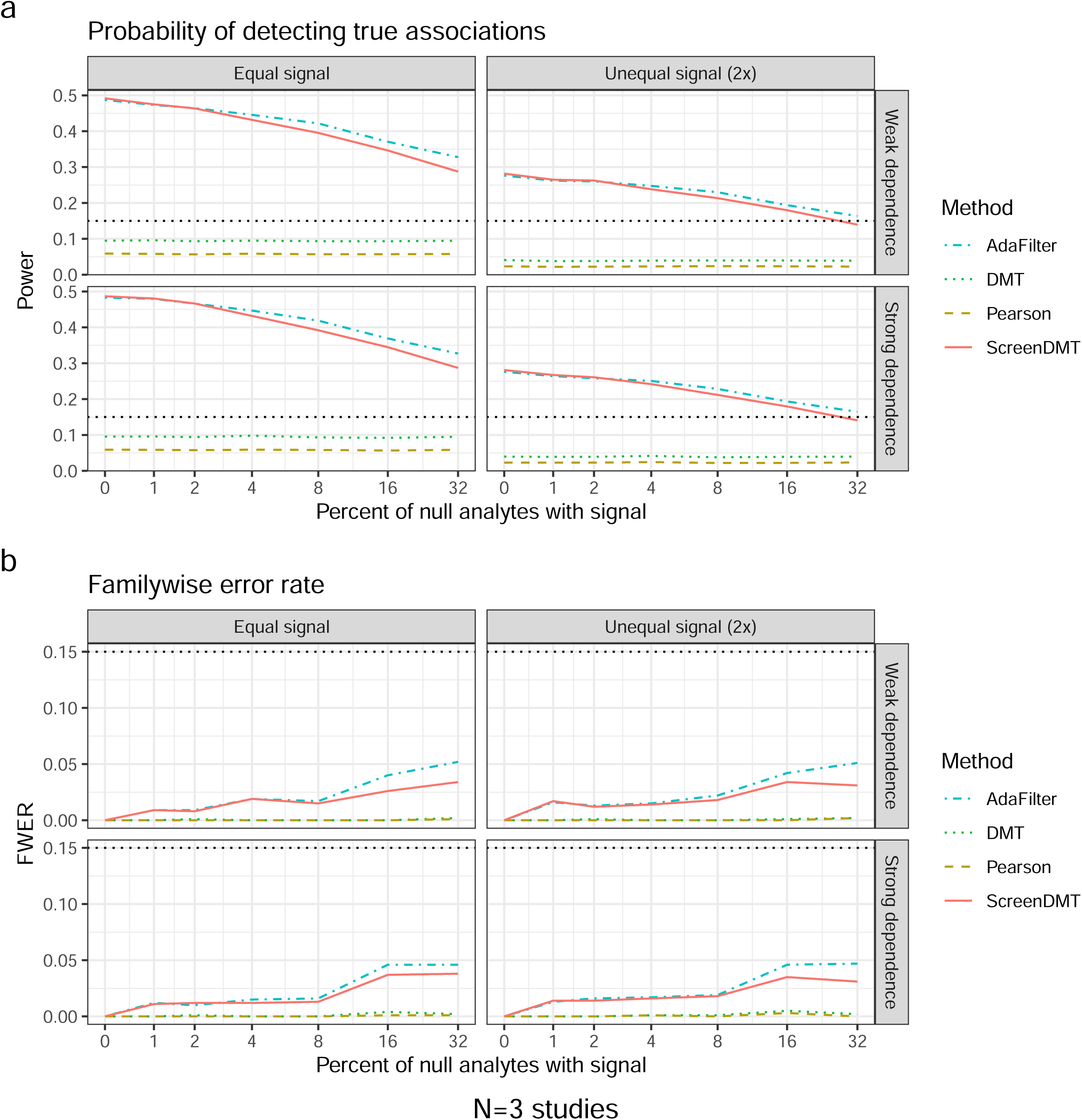
Power and FWER for multiple methods with 3 studies. The x-axis represents the percent of analytes that do not truly directionally replicate across all N studies but show effect in a subset of the studies or show effects of opposite direction in all N studies. Five percent of the analytes truly directionally replicate across all N studies by having only positive or only negative effects. The remaining analytes do not have any true effect in any study. (A) shows the power and (B) shows the family-wise error rate.

### Application across three human cohorts

We applied ScreenDMT to untargeted blood metabolomic profiles of the 228 overlapping metabolites from the Atherosclerosis Risk in Communities Study (ARIC), Multi-Ethnic Study of Atherosclerosis (MESA), and Hispanic Community Health Study/Study of Latinos (HCHS/SOL), which are characterized in Table 1. For comparison, we also applied AdaFilter and Joint Mirror. From each dataset, we separately tested the cross-sectional association of the metabolites to insomnia, long sleep, short sleep, and excessive daytime sleepiness (EDS). We applied the methods to the signed estimates and p-values from each of these metabolites across all three datasets. With ScreenDMT we found two metabolite-outcome pairs that had FDR below our threshold of 15%, as we used previously (Dreyfuss, Djordjilović et al. 2024), which were cortisol with EDS (negative association, DMT z-score = −2.53, DMT p-value = 0.01, ScreenDMT FDR = 0.02) and n-formylmethionine with long sleep (positive association, DMT z-score = 4.31, DMT p-value = 0.04, ScreenDMT FDR = 0.08). AdaFilter only identified cortisol with EDS (at an FDR = 0.09) and Joint Mirror did not identify any replication. All results are given in Table S1.

**Table 1:** Study characteristics.

| Characteristic | HCHS/SOL | ARIC | MESA |
| --- | --- | --- | --- |
| N | 6180 | 1337 | 1874 |
| Age (mean, SD) | 48.28<br>(15.27) | 62.7<br>(5.6) | 68.20<br>(9.04) |
| BMI (mean, SD) | 29.93<br>(6.14) | 28.6<br>(4.9) | 28.66<br>(5.53) |
| Female (%) | 52.8 | 51.2 | 53.3 |
| Current smoker (%) | 22.7 | 10.5 | 7 |
| Diabetes (%) | 18.5 | 10.8 | 19.1 |
| Short Sleep (%) | 9.5 | 7.9 | 19.9 |
| Long Sleep (%) | 25 | 5.7 | 16.4 |
| Insomnia (%) | 30 | 52.5 | 28.9 |
| Excess Daytime Sleepiness (%) | 15 | 29.5 | 14.1 |

In Figure 5, we show the five most significant metabolites per sleep disorder on the y- axis (except short sleep since all of its FDRs > 0.95), negative log10(FDR) on the x- axis, and each point represents one cohort or the ScreenDMT or AdaFilter statistic. AdaFilter’s FDR is stronger than ScreenDMT’s for androsterone glucuronide in EDS, whereas ScreenDMT’s is stronger in 14 out of the 15 cases.

**Figure 5:**
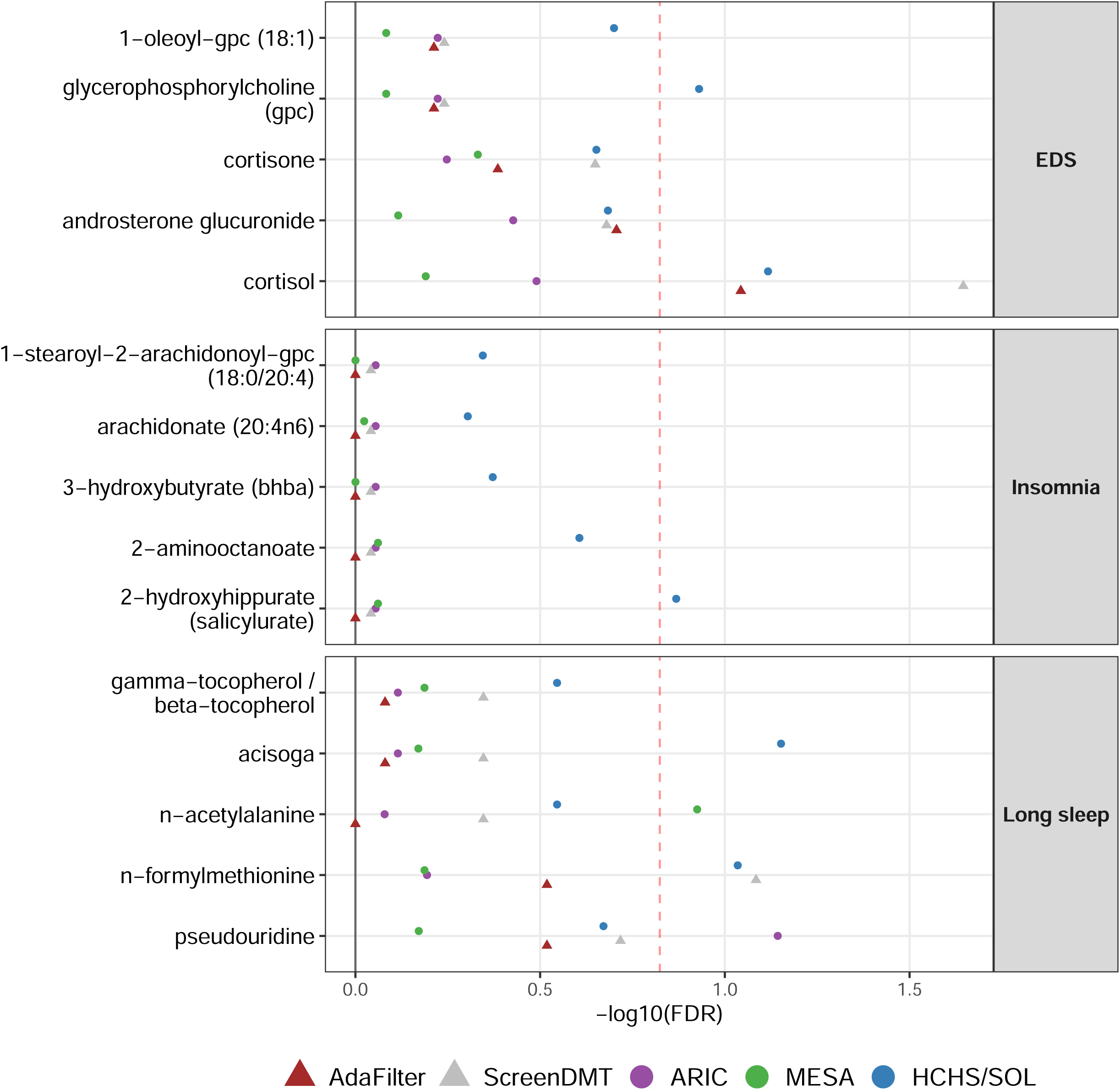
FDRs of the top metabolites per sleep disorder. The five most significant metabolites per sleep disorder are on the y-axis, negative log10(FDR) on the x-axis, and each point represents one cohort or the ScreenDMT or AdaFilter statistic. The dashed line marks FDR = 0.15.

We tested the directional replication of pathways using ScreenDMT, AdaFilter, and Joint Mirror. The results are shown in Figure 6, which is the analog to Figure 5 for pathways (except short sleep since all its pathways had FDR = 1). With ScreenDMT we found the significant pathway-outcome pair of xanthine metabolism with insomnia (negative association, DMT z-score = −2.04, DMT p-value = 0.04, ScreenDMT FDR = 0.12) whose metabolite z-scores are shown in Figure 7, whereas both other methods did not identify any replication. All statistics are in Table S2.

**Figure 6:**
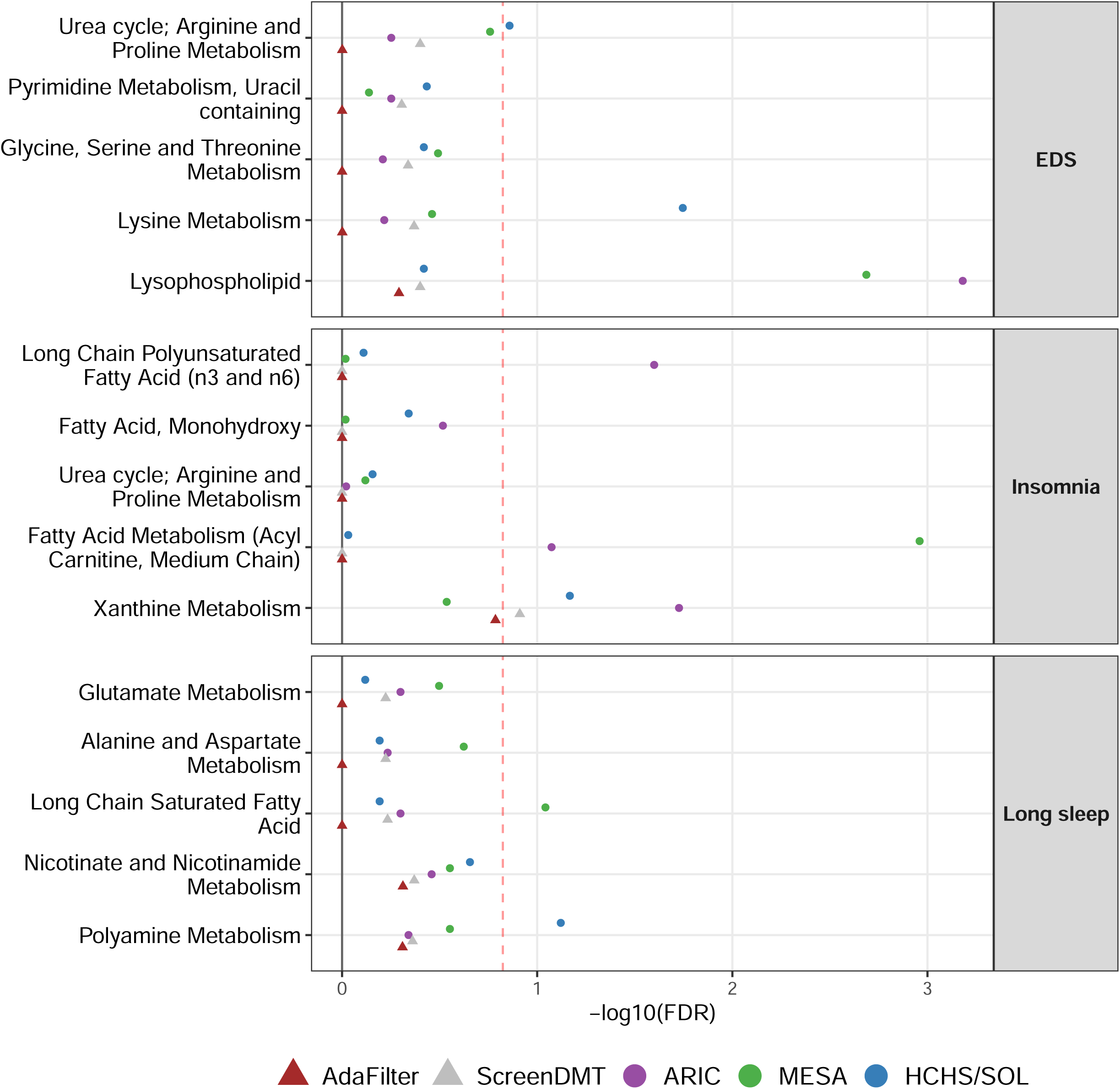
FDRs of the top pathways per sleep disorder. The five most significant pathways per sleep disorder are on the y-axis, negative log10(FDR) on the x-axis, and each point represents one cohort or the ScreenDMT or AdaFilter statistic. The dashed line marks FDR = 0.15.

**Figure 7:**
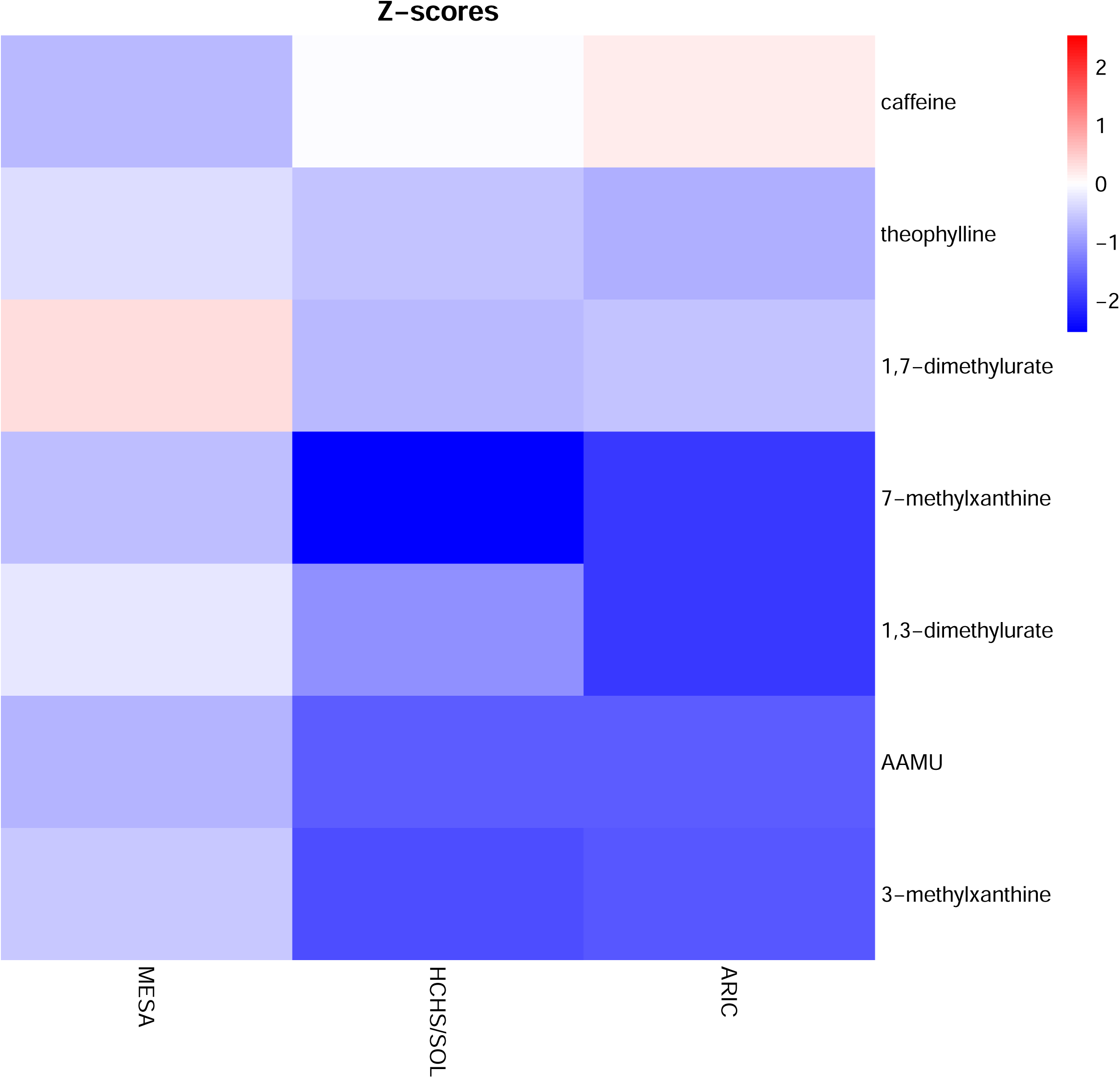
Z-scores of the xanthine metabolism metabolites across cohorts. All metabolites in the Xanthine metabolism that were in our dataset are shown on the y-axis (AAMU = 5-acetylamino-6-amino-3-methyluracil), the cohorts are shown on the x-axis, and the color depicts the z-score.

## Discussion

We extend replication testing with ScreenDMT to an arbitrary number of studies and demonstrate that its primary benefit is for testing replication across three studies. Although this use case may appear to be uncommon, when researchers search for replication across *n* out of *n* studies for *n > 2*, it is most common to use a small *n* such as three (Heller and Yekutieli 2014, Wang, Gui et al. 2022, Liang, Zhang et al. 2025), whereas for larger *n*, it is more common to search for analytes that replicate in at least *r* > 1 out of *n* studies, such as *r=3* out of *n=4* or more.

Our simulations include the recent Joint Mirror procedure (Deng, He et al. 2024), which employs a knockoff-type iterative method that gradually shrinks the rejection region based on progressively revealed information until a conservative estimate of the false discovery proportion is below the target FDR level. This procedure was shown to control the FDR in finite samples and has the property that if the procedure rejects any hypotheses, the number of rejections cannot be less than 1/FDR e.g. if the FDR = 10%, then if it makes any rejections it must reject at least 10 hypotheses, which might explain its conservative results. A procedure we previously included in Dreyfuss et al. (Dreyfuss, Djordjilović et al. 2024) is RepFdr (Heller, Yaacoby et al. 2014), but it did not previously control its error rates with our settings for two studies, and here we found that it also did not control its error rates with our settings for more studies, so we do not show it here.

We found low levels of cortisol to be associated with EDS. Although high levels of pre-sleep cortisol are associated with reduced sleep (Yap, Tung et al. 2024), low levels of post-awakening cortisol are associated with daytime sleepiness (Sørensen, Kåreholt et al. 2024), which is sensible since cortisol promotes wakefulness. We found high levels of n-formylmethionine to be associated with long sleep. This is consistent with a previous study that found that N-formylmethionine is protective for insomnia (Luo, Yin et al. 2025). Lastly, we found low levels of metabolites in xanthine metabolism to be associated with insomnia. Xanthine metabolism’s metabolites in our harmonized data include caffeine and its methylated breakdown products, whose z-scores across the cohorts are shown in Figure 7. Although caffeine consumption late in the day disrupts sleep (Drake, Roehrs et al. 2013, Gardiner, Weakley et al. 2023), our finding is consistent with our previous finding in HCHS/SOL that low levels of hydrocinnamate, a metabolite downstream from caffeic acid found in coffee, are associated with insomnia (Kusters, Santos Cabrera et al. 2026). The associations with coffee metabolites could be due a behavioral adaptation where people with sleeping disorders reduce their caffeine intake late in the day.

The convergence of our replication findings with these independent studies strengthens confidence in both our statistical methodology and the biological relevance of these metabolic pathways in sleep disorders, suggesting these metabolites may serve as biomarkers across diverse populations.

## Methods

### Simulation Studies

We simulated scores of 1,000 metabolites across three studies, and separately we simulated these metabolites across four studies. We simulated each setting 1,000 times and averaged over the simulations. We fixed the proportion of truly directionally replicated (i.e. non-null) metabolites at 0.05. We defined null metabolites as those that do not satisfy the directional replication alternative hypothesis because they are never associated with the outcome, are associated with the outcome in only some of the studies, or are associated with the outcome in all studies but in different directions. Those null metabolites that have some association with the outcome are termed as having some signal.

We varied the proportion of null analytes with some signal across 0, 0.01, 0.02, 0.04, 0.08, 0.16, and 0.32, corresponding to the x-axis in Figures 2-4. All remaining metabolites were simulated with no signal in any study. We evaluated both the setting where truly directionally replicated metabolites had the same magnitude of effect in all studies (the equal-signal setting) and the unequal-signal setting, where these metabolites had different signal magnitudes across studies while preserving the same average signal strength.

In the equal-signal setting, truly directionally replicated metabolites had mean z-scores of 3.5 in every study, with the sign chosen to be either all positive or all negative. In the unequal signal setting, truly directionally replicated metabolites had mean z-scores of 2.33 in half of the studies and 4.67 in the other half, again with a common sign across all studies, so that the average effect magnitude remained 3.5. For the unequal signal setting when the number of studies was odd, one study remained at the baseline magnitude of 3.5. For null analytes with some signal, the study-specific mean z-scores were sampled independently from the set {-3.5, 0, 3.5} with equal probability. If the sampled metabolite corresponded to no signal in all studies or to directional replication across all studies (that is, if all study-specific means were 0, all were positive, or all were negative) then it was discarded and resampled. This produced null analytes with signal patterns that included effects in only a subset of studies as well as effects in opposite directions across studies, while ensuring that they remained under the directional replication null. For each metabolite in each study, statistics were simulated from a normal distribution with standard deviation of one and the specified study-specific mean. Under the per-study null that the mean of the statistic was zero, these statistics were z-scores. From these z-scores we computed two-sided p-values.

### The Hispanic Community Health Study/Study of Latinos

The Hispanic Community Health Study/Study of Latinos (HCHS/SOL) is a community-based prospective cohort study of 16,415 self-identified Hispanic/Latino individuals. The baseline exam took place between 2008 to 2011. Recruitment took place in four field centers located in metropolitan areas (Chicago IL, Miami FL, Bronx NY, and San Diego CA) using a complex survey design, whereby census block unit followed by households were sampled (Lavange, Kalsbeek et al. 2010, Sorlie, Avilés-Santa et al. 2010).

Sleep phenotypes were measured as part of the baseline exam and included sleep duration, insomnia, and excessive daytime sleepiness (EDS). Sleep duration was computed from the questions “what time do you usually go to bed?” and “what time do you usually wake up?” on weekdays and weekends. Short sleep was defined as sleep duration <7 hours, and long sleep was defined as sleep duration >9 hours. Insomnia was based on the Women’s Health Initiative Insomnia Rating Scale (WHIIRS)(Levine, Dailey et al. 2005) with insomnia defined as WHIIRS >9. Excessive daytime sleepiness was based on the Epworth Sleepiness Scale (ESS)(Johns 1991) with EDS defined as ESS >10.

In addition, blood samples were collected for all individuals in a fasting state. Briefly, the extracted serum samples were collected and stored at −80°C. Of these samples, 6,372 were used for metabolomic quantification by Metabolon Inc. (Durham, NC). The samples were sent in two batches first in 2017 (n=4,004) and 2021 (n=2,368).

Individuals in batch 1 were selected at random out of all individuals with available whole genome sequencing at 2017, itself a random sample of HCHS/SOL participants.

Individuals who participated in batch 2 metabolomics assays using baseline study serum samples were selected by a combination of sampling conditions, including a subset of individuals who had estimated glomerular filtration rates in both HCHS/SOL visit 1 and visit 2, and a subset of individuals who participated in the ECHO-SOL ancillary echocardiogram study. Consequently, these batch 2 participants do not represent a random sample from the HCHS/SOL participant population.

Metabolomic quantification was performed using the Discovery HD4 Ultra-High-Performance Liquid Chromatography tandem mass spectrometry (LC-MS) platform. In cases where the same individual had more than one sample (either in the same batch or across batches), one sample was selected at random. Metabolites with 25% or more missing values were excluded. For included metabolites, missing metabolite values were imputed using the half-the-minimum observed value in the sample, under the assumption that their value is below the limit of detection.

For the metabolomics association analyses, metabolite values were rank-normalized separately within each metabolomics batch. The two metabolomics batches were then combined for the primary analysis. We estimated the association between each sleep phenotype and each metabolite using survey-weighted linear regression, with the transformed metabolite level as the outcome and the sleep phenotype as the primary exposure. The complex sampling design of HCHS/SOL was accounted for using sampling weights, strata, and primary sampling units.

The regression model was adjusted for age, sex, study center, and Hispanic/Latino background. Analyses were conducted separately for short sleep, long sleep, EDS, and insomnia. Participants with missing values for the relevant sleep phenotype or model covariates were excluded from the corresponding model. For each sleep phenotype and model, p-values across metabolites were adjusted for multiple testing using the Benjamini–Hochberg FDR.

### Atherosclerosis Risk in Communities Study

The Atherosclerosis Risk in Communities (ARIC) study is a prospective population-based cohort study of 15,792 men and women (Wright, Folsom et al. 2021). Individuals were drawn from four US communities (northwestern suburbs of Minneapolis, Minnesota; Washington County, Maryland; Forsyth County, North Carolina; and Jackson, Mississippi). For the present analysis, 1,337 participants were included who had both metabolomic data at visit 3 (1993-1995), and sleep phenotype data at visit 4 (1996-1998).

In 1996-1998, 1,920 ARIC participants from the Minnesota and Maryland field centers underwent sleep assessments as part of the Sleep Heart Health Study (SHHS)(Quan, Howard et al. 1997). The SHHS Sleep Habits Questionnaire was used to elicit information about habitual sleep duration, insomnia, and excessive daytime sleepiness. To ascertain insomnia, three of the five Women’s Health Initiative Insomnia Rating Scale (WHIIRS)(Levine, Dailey et al. 2005) questions were asked regarding how often individuals experienced (from “0. Never” to “4. Almost always (16-30/month)”) trouble falling asleep, waking up earlier than planned, and trouble getting back to sleep. Responses to these three questions were then summed to create a modified score between 0 and 12, with insomnia defined as a score ≥6. Short sleep was defined as sleep duration <7 hours, and long sleep was defined as sleep duration >9 hours. Excessive daytime sleepiness was based on the Epworth Sleepiness Scale (ESS)(Johns 1991) with EDS defined as ESS >10.

Metabolites from 7,064 ARIC visit 3 participants were measured in two batches: 5,325 samples included through the TOPMed program, and 1,739 more samples outside of TOPMed. For both batches, metabolomic quantification was done by Metabolon Inc. (Durham, NC), using four types of ultra-high-performance liquid chromatography–tandem mass spectroscopy. Compounds were identified using library reference entries of purified standards. Recurrent unknown entities were labeled based on their combination of chromatographic properties and mass spectra. Peaks were quantified using the area under the curve. As the current study spanned multiple days, a data normalization step was performed to correct variation from instrument interday tuning differences. A sample was excluded if it had a metabolite missing rate > 10%, and a metabolite was excluded if it had a missing rate >25%. We then calculated coefficient of variation (CV) using the Bland-Altman method (Bland and Altman 1986) and Spearman correlation coefficient for metabolites, and those that had a CV >50% or Spearman correlation < 0.3 were excluded. For included metabolites, missing metabolite values were imputed using the half-the-minimum observed value in the sample, under the assumption that their value is below the limit of detection.

The metabolomics association analyses in ARIC were conducted using an approach similar to that used in HCHS/SOL, except that standard linear regression models were fit rather than survey-weighted linear regression models, because ARIC does not require incorporation of the HCHS/SOL complex survey design. For each sleep phenotype, we modeled each metabolite as the outcome and the sleep phenotype as the primary exposure. Our regression model adjusted for age, sex, study center, and the difference in days between an individual’s visit 3 and visit 4 exam date.

Participants with missing values for the relevant sleep phenotype or covariates were excluded from the corresponding model. P-values across metabolites were adjusted for multiple testing using the Benjamini–Hochberg FDR.

### The Multi-Ethnic Study of Atherosclerosis

The Multi-Ethnic Study of Atherosclerosis (MESA) is a study of the characteristics of subclinical cardiovascular disease and the risk factors that predict progression to clinically overt cardiovascular disease or progression of the subclinical disease (Bild, Bluemke et al. 2002). MESA consists of a diverse, community-based sample of an initial 6,814 men and women aged 45-84 years without known cardiovascular disease at baseline. Thirty-eight percent of the recruited participants were White, 28 percent African American, 22 percent Hispanic, and 12 percent of Chinese descent. Participants were recruited from six field centers across the United States: Baltimore City and Baltimore County, Maryland; Chicago, Illinois; Forsyth County, North Carolina; Los Angeles County, California; New York, New York; and St. Paul, Minnesota. The first examination took place over two years, from July 2000 to July 2002, and has been followed by additional examinations.

Participants are being followed for identification and characterization of cardiovascular disease events, including acute myocardial infarction and other forms of coronary heart disease (CHD), stroke, and heart failure; for cardiovascular disease interventions; and for mortality. Follow-up telephone interviews with participants or their proxies are attempted at least annually to identify new hospitalizations and diagnoses. MESA staff request information on hospital admissions for any reason, outpatient CVD diagnoses, and death. Death certificate records are obtained from state vital statistics or the National Death Index. MESA staff record International Classification of Disease (ICD) diagnosis codes for all hospitalizations and request medical records for those with ICD codes related to CVD. MESA staff also request outpatient records for potential CVD events. However, not all hospitalizations and/or CVD event records were fully documented.

The study was approved by the Institutional review boards at all participating institutions, and all participants gave written informed consent. In addition, informed consent was obtained for extensive data sharing (dbGaP) and genetic/omic studies, including candidate genes (NHLBI CARe), genome-wide scans (NHLBI SHARe), exome sequencing (NHLBI ESP) and, most recently, the NHLBI TOPMed program.

Sleep measures were ascertained as part of a sleep ancillary study (Chen, Wang et al. 2015), conducted on or slightly after visit 5 (2010-2013) and included sleep duration, insomnia and excessive daytime sleepiness based on the same questions as HCHS/SOL.

MESA metabolomics data were assayed by the Broad Institute – Beth Isreal Deaconess Medical Center platform. Metabolomics data were provided in four files, one for each liquid chromatography – mass spectrometry (LC-MS) method. A drift correction sample follow by a pooled control sample was added after every 10 samples processed in the mass spectrometer. We used the pooled plasma samples to compute the coefficient of variation (CV) for each metabolite. We required CV<0.3 to keep a metabolite in the data. A handful of metabolites were reported in more than one of the LC-MS methods. In this case, we kept the metabolite with the lower CV. Some individuals had repeated samples, in this case we chose one sample at random. Metabolites with 25% or more missing values were excluded. For included metabolites, missing metabolite values were imputed using the half-the-minimum observed value in the sample, under the assumption that their value is below the limit of detection.

The metabolomics association analyses in MESA were conducted using an approach similar to that used in HCHS/SOL, except that standard linear regression models were fit rather than survey-weighted linear regression models, because MESA does not require incorporation of the HCHS/SOL complex survey design. For each sleep phenotype, we modeled each metabolite as the outcome and the sleep phenotype as the primary exposure. The regression model adjusted for age, sex, study center, and self-reported race/ethnicity.

Participants with missing values for the relevant sleep phenotype or covariates were excluded from the corresponding model. P-values across metabolites were adjusted for multiple testing using the Benjamini–Hochberg FDR.

### Harmonizing metabolomics

Metabolite annotations from the ARIC, MESA, and HCHS/SOL cohorts were harmonized to create a unified dataset. First, metabolite nomenclature was standardized across all cohorts by converting names to lowercase and replacing instances of “ic acid” with “ate”. For the MESA cohort, which utilized four distinct LC-MS/MS platforms, internal standards and metabolites lacking names or identifiers were removed. Duplicate annotations across the MESA platforms were systematically resolved by prioritizing entries with the highest database specificity (Level 1 identification); in cases of tied specificity, the annotation with the lowest coefficient of variation (CV) was retained. Because the ARIC and HCHS/SOL cohorts both utilized the Metabolon platform, their datasets were merged directly using shared proprietary compound identifiers after filtering out unnamed or partially characterized molecules.

Finally, the MESA dataset was integrated with the combined ARIC/HCHS/SOL dataset via a two-step matching protocol: exact matching on Human Metabolome Database (HMDB) IDs, followed by matching on cleaned metabolite names (excluding spaces and hyphens) for any remaining unlinked compounds. Duplicate mappings resulting from metabolites with multiple HMDB IDs were subsequently removed to ensure a unique, final alignment across all three cohorts. All steps were manually checked. This resulted in a dataset of 235 metabolites. Since some of these metabolites did not have summary statistics in some cohorts due to missingness, our analysis was based on 228 of these 235 metabolites.

### Replication analysis

We applied the replication methods to the metabolite-level and pathway-level summary statistics. We calculated the pathway-level summary statistics by testing pathways defined by Metabolon (Morrisville, NC) using the Camera method (Wu and Smyth 2012) in the R package Limma (Ritchie, Phipson et al. 2015) in each cohort, yielding signed statistics and p-values of pathway enrichment per cohort. The replication methods we applied were from the R package JMirror version 0.1.0 from https://github.com/denglinsui/JMirror, the R package AdaFilter version 1.0.0 from https://github.com/jingshuw/adaFilter, and the R package DirectionalMaxPTest version 0.0.2 from https://github.com/jdreyf/DirectionalMaxPTest. These were installed using the install_github function from the remotes R package. The replication analysis can be reproduced from https://github.com/jdreyf/ScreendmtMultiAricMesaSol.

## Author contributions

Conceptualization: JMD, TS. Formal analysis: JMD, DD, YZ, MRB. Funding acquisition: TS. Investigation: all authors. Methodology: JMD, VD, TS. Supervision: TS. Writing – Original Draft Preparation: JMD, DD, YZ, MRB, HP, VD, TS. Writing – Review & Editing: all authors.

## Acknowledgements

The work was supported by National Institute on Aging (NIA) grants R01AG080598 and R03AG102038 and National Heart Lung and Blood Institute (NHLBI) grant R01HL161012. Support for metabolomics data was graciously provided by the JLH Foundation (Houston, Texas) and by NHLBI grant R01HL141824.

The authors thank the staff and participants of HCHS/SOL for their important contributions. Investigators website - http://www.cscc.unc.edu/hchs/. The Hispanic Community Health Study/Study of Latinos is a collaborative study supported by contracts from the National Heart, Lung, and Blood Institute (NHLBI) to the University of North Carolina (HHSN268201300001I / N01-HC-65233), University of Miami (HHSN268201300004I / N01-HC-65234), Albert Einstein College of Medicine (HHSN268201300002I / N01-HC-65235), University of Illinois at Chicago (HHSN268201300003I / N01- HC-65236 Northwestern Univ), and San Diego State University (HHSN268201300005I / N01-HC-65237). The following Institutes/Centers/Offices have contributed to the HCHS/SOL through a transfer of funds to the NHLBI: National Institute on Minority Health and Health Disparities, National Institute on Deafness and Other Communication Disorders, National Institute of Dental and Craniofacial Research, National Institute of Diabetes and Digestive and Kidney Diseases, National Institute of Neurological Disorders and Stroke, NIH Institution-Office of Dietary Supplements.

The MESA projects are conducted and supported by the National Heart, Lung, and Blood Institute (NHLBI) in collaboration with MESA investigators. Support for MESA is provided by contracts 75N92025D00022, 75N92020D00001, HHSN268201500003I, N01-HC-95159, 75N92025D00026, 75N92020D00005, N01-HC-95160, 75N92020D00002, N01-HC-95161, 75N92025D00024, 75N92020D00003, N01-HC-95162, 75N92025D00027, 75N92020D00006, N01-HC-95163, 75N92025D00025, 75N92020D00004, N01-HC-95164, 75N92025D00028, 75N92020D00007, N01-HC-95165, N01-HC-95166, N01-HC-95167, N01-HC-95168, N01-HC-95169, UL1-TR-000040, UL1-TR-001079, UL1-TR-001420, UL1TR001881, and R01HL105756. The authors thank the MESA participants and the MESA investigators and staff for their valuable contributions. A full list of participating MESA investigators and institutions can be found at http://www.mesa-nhlbi.org. We gratefully acknowledge the studies and participants who provided biological samples and data for TOPMed. Molecular data for the Trans-Omics in Precision Medicine (TOPMed) program was supported by the NHLBI. Metabolomics for “NHLBI TOPMed: Multi-Ethnic Study of Atherosclerosis (MESA)” (phs001416) was performed at Broad Institute and Beth Israel Metabolomics Platform (HHSN268201600038I). Core support including centralized genomic read mapping and genotype calling, along with variant quality metrics and filtering were provided by the TOPMed Informatics Research Center (3R01HL-117626-02S1; contract HHSN268201800002I). Core support including phenotype harmonization, data management, sample-identity QC, and general program coordination were provided by the TOPMed Data Coordinating Center (R01HL-120393; U01HL120393; contract HHSN268201800001I).

The Atherosclerosis Risk in Communities study has been funded in whole or in part with Federal funds from the National Heart, Lung, and Blood Institute, National Institutes of Health, Department of Health, and Human Services, under Contract nos. (75N92022D00001, 75N92022D00002, 75N92022D00003, 75N92022D00004, 75N92022D00005). The authors thank the staff and participants of the ARIC study for their important contributions. Visit 3 metabolomics measurements were sponsored in by the National Heart, Lung, and Blood Institute (NHLBI) grant R01HL141824 and Trans-Omics in Precision Medicine (TOPMed) program. Metabolomics for “NHLBI TOPMed - NHGRI CCDG: Atherosclerosis Risk in Communities (ARIC)” (phs001211) was performed at the University of Texas Health Science Center at Houston/Metabolon (HHSN268201600033I).

## Data availability statement

In accordance with participants informed consent, MESA, HCHS/SOL and ARIC data are available through data use agreement in dbGaP according to the study specific accessions. MESA phenotypes are available in: phs000209; MESA metabolomics data have been deposited and will become available in: phs001416; and HCHS/SOL phenotypes: phs000810. HCHS/SOL metabolomics data are available via data use agreement with the HCHS/SOL Data Coordinating Center at the University of North Carolina at Chapel Hill, see collaborators website: https://sites.cscc.unc.edu/hchs/. The ARIC phenotypes are available in: phs000280.v8.p2 or BioLINCC (https://biolincc.nhlbi.nih.gov/studies/aric/). The ARIC metabolomic data can be requested through the study’s Data Coordinating Center upon an approved manuscript proposal and Data and Materials Distribution Agreement (DMDA). All data generated or analyzed during this study are included in this published article and its supplementary information file.

The full replication analysis from the metabolite-level summary statistics and harmonized annotation can be reproduced from https://github.com/jdreyf/ScreendmtMultiAricMesaSol. The ScreenDMT method is available from https://github.com/jdreyf/DirectionalMaxPTest.

## Supporting information

Table S1 Metabolite-level statistics per cohort and replication results with AdaFilter and ScreenDMT.

Table S2 Pathway-level statistics per cohort and replication results with AdaFilter and ScreenDMT.

### Note S1

Statement of the proposition and theorem and their mathematical proofs.

