## Supplementary material for "ScreenDMT reveals metabolites replicably associated with sleep disorders across three human cohorts": Note S1

### Note S1: ScreenDMT for multiple replication procedure and validation

#### Problem statement

Let $\boldsymbol{\theta} = [\theta_1, \theta_2, ..., \theta_n]$ be a vector of finite, real-valued parameters. The $\theta_i$'s represent parameters pertaining to the same biological entity measured across $n \geq 2$ studies, $n \in \mathbb{N}$. Each study has sample size $\nu_i$. We consider the problem of detecting replication in the same direction across studies. We define a parameter regarding the sign,

$$S = \begin{cases} 1, & \text{if } \boldsymbol{\theta} \succ 0 \text{ or } \boldsymbol{\theta} \prec 0 \\ 0, & \text{otherwise} \end{cases} \tag{1}$$

where $\boldsymbol{\theta} \succ 0$ denotes that all elements of the vector are positive. Then we are interested in testing the directional null hypothesis:

$$H_0 : S = 0 \quad \text{against} \quad \bar{H}_0 : S = 1.$$

Let $\mathcal{I} = \{1, 2, ..., n\}$ and $\mathcal{I} \setminus k = \{j \in \mathcal{I} | j \neq k\}$ for $k \in \mathcal{I}$. For $i \in \mathcal{I}$, we assume that we have $\sqrt{\nu}_i$-consistent and asymptotically normal estimates $\hat{\theta}_i$ of $\theta_i$ such that

$$\sqrt{\nu}_i(\hat{\theta}_i - \theta_i) \xrightarrow{d} N(0, \sigma_i^2)$$

where the convergence is in distribution and $0 < \sigma_i < \infty$, where $\sigma_i$ is known. Then we can define component $p$-values $P_1, P_2, ..., P_n$ for testing $H_1 : \theta_1 = 0$, $H_2 : \theta_2 = 0$, $\ldots$, $H_n : \theta_n = 0$ respectively, as

$$P_i = 2\Phi\left(-\frac{\sqrt{\nu}_i|\hat{\theta}_i|}{\sigma_i}\right) \tag{2}$$

where $\Phi(\cdot)$ is the standard normal cumulative distribution function and $\boldsymbol{P} = [P_1, P_2, ..., P_n]$. The procedures below are valid for large samples if consistent estimates of $\sigma_i$ are used instead. We further assume that the component estimates of $\hat{\boldsymbol{\theta}} = [\hat{\theta}_1, \hat{\theta}_2, ..., \hat{\theta}_n]$ are mutually independent, a plausible assumption for replication.

To simplify notation we let

$$\delta_i = \frac{\theta_i \sqrt{\nu}_i}{\sigma_i}$$

so that Equation 2 becomes $P_i = 2\Phi\left(-|\hat{\delta}_i|\right)$ and $\hat{\delta}_i - \delta_i \sim N(0, 1)$. This is without loss of generality (WLOG), since in any dataset $\sqrt{\nu}_i/\sigma_i$ is a constant coefficient of $\hat{\theta}_i$. The constant is always positive, so $\delta_i$ could replace $\theta_i$ in the problem statement, where the analog of $\boldsymbol{\theta}$ is $\boldsymbol{\delta} = [\delta_1, \delta_2, ..., \delta_n]$. We assume $\theta_i$ and $\sigma_i$ are fixed, so $|\delta_i| \to \infty$ if and only if $\theta_i \neq 0$ and $\nu_i \to \infty$.

### MaxP test

The non-directional (ND) null hypothesis for replication of $n$ studies is

$$H_0 : H_0^{ND} = \cup_{i \in \mathcal{I}} H_i \quad \text{against} \quad \bar{H}_0^{ND} : \cap_{i \in \mathcal{I}} \bar{H}_i.$$

For this hypothesis, a common approach is the MaxP or joint significance test, which has p-value $\bar{P} = \max\{P_1, P_2, ..., P_n\}$. It does not account for direction. For the case of $n = 2$ studies, Liu et al. (2022) showed that the MaxP test was the likelihood ratio test (LRT). Liu et al. (2022) were actually examining the case of $n = 2$ in the context of causal mediation analysis. The reason that the mediation null is equivalent to the replication null with $n = 2$ is because mediation also tests for joint significance, of the exposure-mediator and mediator-outcome effects. We extend that proof for general $n \geq 2$.

#### Proposition: MaxP test for $n \geq 2$ is the LRT for the non-directional hypothesis

#### Proof

Consider applying a LRT based on the asymptotic sampling distribution of $\hat{\boldsymbol{\theta}}$. The LRT rejects $H_0^{ND}$ for large values of:

$$\tau = -2 \ln \left[ \frac{\max_{\boldsymbol{x} \in \Theta_0^{ND}} \prod_{i=1}^n \phi\{\sqrt{\nu_i}(\hat{\theta}_i - x_i)/\sigma_i\}}{\max_{\boldsymbol{y} \in \mathbb{R}^n} \prod_{i=1}^n \phi\{\sqrt{\nu_i}(\hat{\theta}_i - y_i)/\sigma_i\}} \right]$$

where $\Theta_0^{ND} = \{\boldsymbol{u} : \boldsymbol{u} \in H_0^{ND}\}$ is the null region of the parameter space ($\bar{\Theta}_0^{ND}$ is the alternative region), $\phi(\cdot)$ is the standard normal density function, and $\boldsymbol{x}$ and $\boldsymbol{y}$ are vectors of size $n$. The denominator is maximized at $y_i^\star = \hat{\theta}_i$ for all $i = 1, ..., n$, where it becomes a term only involving $\boldsymbol{\sigma}$ that is also in the numerator, so they cancel out. Then,

$$\begin{aligned} \tau &= -2 \max_{\boldsymbol{x} \in \Theta_0^{ND}} \sum_{i=1}^n -\frac{\nu_i}{2} \left( \frac{\hat{\theta}_i - x_i}{\sigma_i} \right)^2 \\ &= \min_{\boldsymbol{x} \in \Theta_0^{ND}} \sum_{i=1}^n \nu_i \left( \frac{\hat{\theta}_i - x_i}{\sigma_i} \right)^2 \end{aligned}$$

If $\hat{\boldsymbol{\theta}} \in \Theta_0^{ND}$, then we will have $\boldsymbol{x}^\star = \hat{\boldsymbol{\theta}}$, yielding $\tau = 0$.

Next, we consider points under the alternative hypothesis, $\hat{\boldsymbol{\theta}} \in \bar{\Theta}_0^{ND}$. The closest $\boldsymbol{x} \in \Theta_0^{ND}$ can be to $\hat{\boldsymbol{\theta}}$ is for $x_i = 0$ for one $i$, $i = 1, ..., n$, whereas for the others $x_i = \hat{\theta}_i$. So the minimization of the sum will select the study with the smallest $\hat{\delta}_i^2$:

$$\tau = \min_{i=1,...,n} \hat{\delta}_i^2$$

and the $p$-value of the LRT is computed as $\mathbb{P}(\chi_1^2 > \tau)$, which is equivalent to

$$\bar{P} = \max_{i=1,...,n} \mathbb{P}(\chi_1^2 > \hat{\delta}_i^2).$$

### Directional MaxP test (DMT)

Let

$$\hat{S} = \begin{cases} 1, & \text{if } \hat{\boldsymbol{\theta}} \succ 0 \text{ or } \hat{\boldsymbol{\theta}} \prec 0 \\ 0, & \text{otherwise.} \end{cases} \tag{3}$$

The DMT p-value, as shown in the main text, is

$$P_{DMT} = \begin{cases} \max\{P_1, P_2, ..., P_n\}/2, & \text{if } \hat{S} = 1 \\ 1, & \text{otherwise.} \end{cases}$$

### ScreenDMT: multiple testing of directional hypotheses with screening

When considering $n \geq 2$ studies with multiple hypotheses (e.g. for many analytes, lipids, genes), we propose the following two-step multiple testing procedure, which generalizes the ScreenDMT procedure we previously validated for two studies (Dreyfuss et al. 2024). The ScreenDMT procedure includes an additional step beyond the DMT based on AdaFilter (Wang et al. 2022) to calculate the adjusted p-values. This step is based on the ordered p-values $P_{(1)} \leq P_{(2)} \leq \cdots \leq P_{(n)}$.

1. Apply the directional MaxP test per analyte to obtain $P_{DMT}$.
2. Calculate adjusted p-values using AdaFilter with filtering p-value $\Lambda = P_{(n-1)}$ and selection p-value $\Xi = \max\{P_{DMT}, P_{(n-1)}\}$. This can be used for calculating adjusted p-values that control the FDR or the FWER.

Dreyfuss et al. (2021) provided an option in step 1 to calculate valid p-values of these parameters in omics data using the R package Limma (Ritchie et al. 2015), which has been found to improve power across omics datasets. This method or any other that calculates valid p-values can be used.

The key property that AdaFilter relies on is a form of conditional validity (Wang et al. 2022). Assuming that for each analyte $\{P_i\}_1^n$ are independent among themselves, the property states that when $H_0$ is true:

$$\mathbb{P}(\Xi < u | \Lambda < u) \leq u$$

holds for any fixed u > 0 whenever $\mathbb{P}(\Lambda < u) > 0$.

#### Theorem: ScreenDMT for $n \geq 2$ satisfies conditional validity in asymptotic sample sizes

#### Proof

To demonstrate conditional validity under $H_0$ when $\nu_i \to \infty$ for all $i = 1, ..., n$, we show that this holds on the boundary of the parameter space where $\theta_1 = 0 \cup \theta_2 = 0 \cup ... \cup \theta_n = 0$ and off the boundary, where we show that the gradient is zero. We have,

$$
\begin{aligned}
\mathbb{P}(\Xi < u | \Lambda < u) =& \mathbb{P}(\max\{P_{DMT}, P_{(n-1)}\} < u | P_{(n-1)} < u) \\
=& \mathbb{P}(P_{DMT} < u, P_{(n-1)} < u | P_{(n-1)} < u) \\
=& \mathbb{P}(P_{DMT} < u, P_{(n-1)} < u)/\mathbb{P}(P_{(n-1)} < u) \\
=& \mathbb{P}(P_{DMT} < u | P_{(n-1)} < u)
\end{aligned}
$$

so we consider $\mathbb{P}(P_{DMT} < u | P_{(n-1)} < u)$ below. For $u \geq 1$, we trivially have $\mathbb{P}(P_{DMT} < u | P_{(n-1)} < u) \leq u$. For $0 < u < 1$, using the law of total probability, we continue

$$
\begin{aligned}
=& \mathbb{P}(P_{DMT} < u, \hat{S} = 1 | P_{(n-1)} < u) + \mathbb{P}(P_{DMT} < u, \hat{S} = 0 | P_{(n-1)} < u) \\
=& \mathbb{P}(P_{DMT} < u, \hat{S} = 1 | P_{(n-1)} < u), \text{ since } \hat{S} = 0 \implies P_{DMT} = 1, \\
=& \mathbb{P}(P_{(n)} < 2u, \hat{S} = 1 | P_{(n-1)} < u) \\
=& \mathbb{P}(\hat{S} = 1)\mathbb{P}(P_{(n)} < 2u | P_{(n-1)} < u), \text{ as } \hat{S} \perp\!\!\!\perp \boldsymbol{P},
\end{aligned}
\tag{4}
$$

where $\hat{S} \perp\!\!\!\perp \boldsymbol{P}$ because each $P_i$ is calculated based on $|\hat{\theta}_i|$, which is independent of $\hat{S}$.

We have previously shown that under $H_0$, $\mathbb{P}(\hat{S} = 1) \leq \frac{1}{2}$ when there are two studies (Dreyfuss et al. 2021). Although that proof was in the context of mediation, it is equivalent to replication with two studies. We now extend this upper bound on $\mathbb{P}(\hat{S} = 1)$ to $n \geq 2$ studies, since we will use this upper bound to show conditional validity.

We proceed by induction. Let $\hat{S}$ in the context of $n$ studies be represented as $\hat{S}_n$. The base case is that $\mathbb{P}(\hat{S}_2 = 1) \leq \frac{1}{2}$, which we previously proved. For the inductive step, let $\mathbb{P}(\hat{S}_k = 1) \leq \frac{1}{2}$ for some $k \geq 2, k \in \mathbb{N}$. Then,

$$
\begin{aligned}
\mathbb{P}(\hat{S}_{k+1} = 1) =& \mathbb{P}(\hat{S}_k = 1, \text{sign}(\hat{\delta}_k) = \text{sign}(\hat{\delta}_{k+1})) \\
\leq& \mathbb{P}(\hat{S}_k = 1) \\
\leq& \frac{1}{2}
\end{aligned}
$$

which shows that $\mathbb{P}(\hat{S}_n = 1) \leq \frac{1}{2}$ for all $n \geq 2$.

For $u > \frac{1}{2}$ we know that $\boldsymbol{P} \prec 2u$, so Equation 4 becomes

$$
\begin{aligned}
\mathbb{P}(P_{DMT} < u | P_{(n-1)} < u) &= \mathbb{P}(\hat{S} = 1)\mathbb{P}(P_{(n)} < 2u | P_{(n-1)} < u) \\
&= \mathbb{P}(\hat{S} = 1) \\
&\leq \frac{1}{2} \\
&< u.
\end{aligned}
$$

We now consider $\mathbb{P}(P_{DMT} < u | P_{(n-1)} < u)$ under $H_0$ for $0 < u \leq \frac{1}{2}$.

$$
\mathbb{P}(P_{(n)} < 2u, \hat{S} = 1 | P_{(n-1)} < u) = \frac{\mathbb{P}(P_{(n)} < 2u, P_{(n-1)} < u, \hat{S} = 1)}{\mathbb{P}(P_{(n-1)} < u)} \tag{5}
$$

We would like to represent Equation 5 in terms of $\boldsymbol{\delta}$, so we can evaluate it under the null. We begin with its denominator. We define the event $P_{(n-1)} < u$ in terms of unordered p-values, which can be defined in terms of $\boldsymbol{\delta}$. We write the event $P_{(n-1)} < u$ as $P_i < u$ for all $i \in \mathcal{I} \setminus k$ for a single $k \in \mathcal{I}$. So we can write the denominator as the probability of the union of these events for each $k \in \mathcal{I}$.

We calculate this probability for all $k$ according to the inclusion-exclusion principle. Given our definition of an event, observe that the intersection of two or more of these events resolves to the event that $P_i < u$ for all $i \in \mathcal{I}$. Then we find that,

$$\begin{aligned}
\mathbb{P}(P_{(n-1)} < u) =& \mathbb{P}\left(\bigcup_{i=1}^{n}\left(\bigcap_{j\in\mathcal{I}\setminus i} P_j < u\right)\right) \\
=& \sum_{i=1}^{n}\mathbb{P}\left(\bigcap_{j\in\mathcal{I}\setminus i} P_j < u\right) - \binom{n}{2}\mathbb{P}\left(\bigcap_{i=1}^{n} P_i < u\right) + \binom{n}{3}\mathbb{P}\left(\bigcap_{i=1}^{n} P_i < u\right) - ... \\
=& \sum_{i=1}^{n}\mathbb{P}\left(\bigcap_{j\in\mathcal{I}\setminus i} P_j < u\right) + \psi(n)\mathbb{P}\left(\bigcap_{i=1}^{n} P_i < u\right)
\end{aligned}$$

where $\psi(n) = \sum_{k=2}^{n}(-1)^{k-1}\binom{n}{k}$ for $n \geq 2$. We can simplify this to $\psi(n) = 1 - n$ using the binomial theorem.

We expand the probability of all $P_i < u$ in terms of $\hat{\boldsymbol{\delta}}$ using the definition of $P_i$ and that $\{P_i\}_1^n$ are independent among themselves.

$$\begin{aligned}
\mathbb{P}\left(\bigcap_{i=1}^{n} P_i < u\right) =& \prod_{i=1}^{n}\mathbb{P}(2\Phi(-|\hat{\delta}_i|) < u) \\
=& \prod_{i=1}^{n}\mathbb{P}(|\hat{\delta}_i| > -\Phi^{-1}(u/2)) \\
=& \prod_{i=1}^{n}\left[\mathbb{P}(\hat{\delta}_i > -\Phi^{-1}(u/2)) + \mathbb{P}(-\hat{\delta}_i > -\Phi^{-1}(u/2))\right] \\
=& \prod_{i=1}^{n}\left[\mathbb{P}(-\hat{\delta}_i + \delta_i < \Phi^{-1}(u/2) + \delta_i) + \mathbb{P}(\hat{\delta}_i - \delta_i \leq \Phi^{-1}(u/2) - \delta_i)\right] \\
=& \prod_{i=1}^{n}\left[\Phi(\Phi^{-1}(u/2) + \delta_i) + \Phi(\Phi^{-1}(u/2) - \delta_i)\right]
\end{aligned}$$

Following this reasoning, we can also derive

$$\mathbb{P}\left(\bigcap_{j\in\mathcal{I}\setminus i} P_j < u\right) = \prod_{j\in\mathcal{I}\setminus i}[\Phi(\Phi^{-1}(u/2) + \delta_j) + \Phi(\Phi^{-1}(u/2) - \delta_j)].$$

Putting these together to resolve $\mathbb{P}(P_{(n-1)} < u)$ in terms of $\hat{\boldsymbol{\delta}}$, we have

$$\mathbb{P}(P_{(n-1)} < u) = \sum_{i=1}^{n}\prod_{j\in\mathcal{I}\setminus i}[\Phi(\Phi^{-1}(u/2)+\delta_j)+\Phi(\Phi^{-1}(u/2)-\delta_j)]+(1-n)\prod_{i=1}^{n}[\Phi(\Phi^{-1}(u/2)+\delta_i)+\Phi(\Phi^{-1}(u/2)-\delta_i)].$$

We now turn to its numerator of Equation 5.

$$
\begin{aligned}
&\mathbb{P}(P_{(n)} < 2u, P_{(n-1)} < u, \hat{S} = 1) \\
=&\mathbb{P}\left(\bigcup_{i=1}^{n}\left(P_i < 2u, \hat{S} = 1, \bigcap_{j\in\mathcal{I}\setminus i} P_j < u\right)\right) \\
=&\sum_{i=1}^{n}\mathbb{P}\left(P_i < 2u, \hat{S} = 1, \bigcap_{j\in\mathcal{I}\setminus i} P_j < u\right) + \left[-\binom{n}{2} + \binom{n}{3} - \binom{n}{4} + ...\binom{n}{n}\right]\mathbb{P}\left(\hat{S} = 1, \bigcap_{j=1}^{n} P_j < u\right) \\
=&\sum_{i=1}^{n}\mathbb{P}\left(P_i < 2u, \hat{S} = 1, \bigcap_{j\in\mathcal{I}\setminus i} P_j < u\right) + \psi(n)\mathbb{P}\left(\hat{S} = 1, \bigcap_{j=1}^{n} P_j < u\right) \\
=&\sum_{i=1}^{n}\mathbb{P}\left(P_i < 2u, \hat{S} = 1, \bigcap_{j\in\mathcal{I}\setminus i} P_j < u\right) + (1-n)\mathbb{P}\left(\hat{S} = 1, \bigcap_{j=1}^{n} P_j < u\right)
\end{aligned}
$$

We evaluate the first term for a single $i$, arbitrarily choosing $i = 1$,

$$
\begin{aligned}
&\mathbb{P}\left(P_1 < 2u, \hat{S} = 1, \bigcap_{j=2}^{n} P_j < u\right) \\
=&\mathbb{P}(2\Phi(-|\hat{\delta}_1|) < 2u, 2\Phi(-|\hat{\delta}_2|) < u, ..., 2\Phi(-|\hat{\delta}_n|) < u, \hat{S} = 1) \\
=&\mathbb{P}(-|\hat{\delta}_1| < \Phi^{-1}(u), -|\hat{\delta}_2| < \Phi^{-1}(u/2), ..., -|\hat{\delta}_n| < \Phi^{-1}(u/2), \hat{S} = 1) \\
=&\mathbb{P}(|\hat{\delta}_1| > -\Phi^{-1}(u), |\hat{\delta}_2| > -\Phi^{-1}(u/2), ..., |\hat{\delta}_n| > -\Phi^{-1}(u/2), \hat{S} = 1) \\
=&\mathbb{P}(\hat{\delta}_1 > -\Phi^{-1}(u), \hat{\delta}_2 > -\Phi^{-1}(u/2), ..., \hat{\delta}_n > -\Phi^{-1}(u/2))+ \\
&\mathbb{P}(-\hat{\delta}_1 > -\Phi^{-1}(u), -\hat{\delta}_2 > -\Phi^{-1}(u/2), ..., -\hat{\delta}_n > -\Phi^{-1}(u/2)), \text{ since } \hat{\boldsymbol{\delta}} \succ 0 \text{ or } \hat{\boldsymbol{\delta}} \prec 0, \\
=&\mathbb{P}(-\hat{\delta}_1 + \delta_1 < \Phi^{-1}(u) + \delta_1)\prod_{j=2}^{n}\mathbb{P}(-\hat{\delta}_j + \delta_j < \Phi^{-1}(u/2) + \delta_j)+ \\
&\mathbb{P}(\hat{\delta}_1 - \delta_1 < \Phi^{-1}(u) - \delta_1)\prod_{j=2}^{n}\mathbb{P}(\hat{\delta}_j - \delta_j < \Phi^{-1}(u/2) - \delta_j) \\
=&\Phi(\Phi^{-1}(u) + \delta_1)\prod_{j=2}^{n}\Phi(\Phi^{-1}(u/2) + \delta_j) + \Phi(\Phi^{-1}(u) - \delta_1)\prod_{j=2}^{n}\Phi(\Phi^{-1}(u/2) - \delta_j).
\end{aligned}
$$

Similarly for the second term,

$$
\begin{aligned}
\mathbb{P}\left(\hat{S} = 1, \bigcap_{j=1}^{n} P_j < u\right) =&\mathbb{P}(\hat{\delta}_1 > -\Phi^{-1}(u/2), \hat{\delta}_2 > -\Phi^{-1}(u/2), ..., \hat{\delta}_n > -\Phi^{-1}(u/2))+ \\
&\mathbb{P}(-\hat{\delta}_1 > -\Phi^{-1}(u/2), -\hat{\delta}_2 > -\Phi^{-1}(u/2), ..., -\hat{\delta}_n > -\Phi^{-1}(u/2)) \\
=&\prod_{j=1}^{n}\mathbb{P}(-\hat{\delta}_j + \delta_j < \Phi^{-1}(u/2) + \delta_j) + \prod_{j=1}^{n}\mathbb{P}(\hat{\delta}_j - \delta_j < \Phi^{-1}(u/2) - \delta_j) \\
=&\prod_{j=1}^{n}\Phi(\Phi^{-1}(u/2) + \delta_j) + \prod_{j=1}^{n}\Phi(\Phi^{-1}(u/2) - \delta_j).
\end{aligned}
$$

Putting all the terms in Equation 5 together under $H_0$ with $0 < u \leq \frac{1}{2}$,

$$
\begin{aligned}
&\mathbb{P}(P_{DMT} < u | P_{(n-1)} < u) \\
=&\Bigg\{ \sum_{i=1}^{n} \Bigg[ \Phi(\Phi^{-1}(u) + \delta_i) \prod_{j \in \mathcal{I} \setminus i} \Phi(\Phi^{-1}(u/2) + \delta_j) + \Phi(\Phi^{-1}(u) - \delta_i) \prod_{j \in \mathcal{I} \setminus i} \Phi(\Phi^{-1}(u/2) - \delta_j) \Bigg] + \\
&(1-n) \left[ \prod_{i=1}^{n} \Phi(\Phi^{-1}(u/2) + \delta_i) + \prod_{i=1}^{n} \Phi(\Phi^{-1}(u/2) - \delta_i) \right] \Bigg\} / \\
&\Bigg\{ \sum_{i=1}^{n} \prod_{j \in \mathcal{I} \setminus i} \left[ \Phi(\Phi^{-1}(u/2) + \delta_j) + \Phi(\Phi^{-1}(u/2) - \delta_j) \right] + \\
&(1-n) \prod_{i=1}^{n} \left[ \Phi(\Phi^{-1}(u/2) + \delta_i) + \Phi(\Phi^{-1}(u/2) - \delta_i) \right] \Bigg\}
\end{aligned} \tag{6}
$$

**Parameters at origin**

At the origin where $\boldsymbol{\delta} = \mathbf{0}$, Equation 6 is

$$
\begin{aligned}
&\mathbb{P}(P_{DMT} < u | P_{(n-1)} < u, \boldsymbol{\delta} = \mathbf{0}) \\
=&\left\{ \sum_{i=1}^{n} \left[ u \prod_{j \in \mathcal{I} \setminus i} \frac{u}{2} + u \prod_{j \in \mathcal{I} \setminus i} \frac{u}{2} \right] + (1-n) \left[ \prod_{i=1}^{n} \frac{u}{2} + \prod_{i=1}^{n} \frac{u}{2} \right] \right\} / \left\{ \sum_{i=1}^{n} \prod_{j \in \mathcal{I} \setminus i} u + (1-n) \prod_{i=1}^{n} u \right\} \\
=&2 \left\{ nu \left( \frac{u}{2} \right)^{n-1} + (1-n) \left( \frac{u}{2} \right)^{n} \right\} / \left\{ nu^{n-1} + (1-n)u^n \right\} \\
=&u 2^{1-n} \frac{n+1}{n + (1-n)u}
\end{aligned} \tag{7}
$$

for $n \geq 2$. This function's upper bound can be easily obtained because it is a convex function in $u$. We show its convexity by finding its second derivative.

$$
\begin{aligned}
\frac{\partial \mathbb{P}(P_{DMT} < u | P_{(n-1)} < u, \boldsymbol{\delta} = \mathbf{0})}{\partial u} =& 2^{1-n}(n+1) \left\{ \frac{1}{n + (1-n)u} - \frac{(1-n)u}{[n + (1-n)u]^2} \right\} \\
=& \frac{2^{1-n}(n+1)n}{[n + (1-n)u]^2}
\end{aligned}
$$

so

$$
\begin{aligned}
\frac{\partial^2 \mathbb{P}(P_{DMT} < u | P_{(n-1)} < u, \boldsymbol{\delta} = \mathbf{0})}{\partial u^2} =& -2 \frac{2^{1-n}(n+1)n(1-n)}{[n + (1-n)u]^3} \\
=& \frac{2^{2-n}(n+1)n(n-1)}{[n + (1-n)u]^3},
\end{aligned}
$$

which is positive for all $n \geq 2$ and $0 < u \leq \frac{1}{2}$. Because $\mathbb{P}(P_{DMT} < u | P_{(n-1)} < u, \boldsymbol{\delta} = \mathbf{0})$ is convex, we also know that $\mathbb{P}(P_{DMT} < u | P_{(n-1)} < u, \boldsymbol{\delta} = \mathbf{0}) - u$ is convex, so we only need to check the endpoints to see if $\mathbb{P}(P_{DMT} < u | P_{(n-1)} < u, \boldsymbol{\delta} = \mathbf{0}) \leq u$. Therefore we evaluate $u 2^{1-n} \frac{n+1}{n+(1-n)u}$ at $u = 0$, where it is zero, and at $u = \frac{1}{2}$, where it is

$$
2^{-n} \frac{n+1}{n + 0.5(1-n)} = 2^{-n} \frac{n+1}{0.5(n+1)} = 2^{1-n} \leq \frac{1}{2}.
$$

Thus, conditional validity is maintained at the origin for $n \geq 2$.

**Parameters non-negative**

Another case under $H_0$ on the boundary but apart from the origin is where $\delta_i = 0$ for at least one $i \in \mathcal{I}$ but not all $i$ and $\boldsymbol{\delta} \succeq 0$ or $\boldsymbol{\delta} \preceq 0$. Due to the symmetry of this problem, WLOG we can choose to evaluate only $\boldsymbol{\delta} \succeq 0$ with equality for at least one element of the vector but not for all elements. We know what happens when $\delta_i = 0$ from the case above, which holds whether the sample size $\nu_i$ is finite or grows without bound. Recall from the problem statement that in asymptotic sample sizes, $\delta_i > 0$ implies $\delta_i \to \infty$. Then $\Phi(\Phi^{-1}(x) + \delta_i) \to 1$ and $\Phi(\Phi^{-1}(x) - \delta_i) \to 0$ for $0 < x \leq 0.5$.

To represent this case, let $\delta_i = 0$ for all $i \in \mathcal{K} = \{1, 2, ..., K\}$, where $1 \leq K < n$, and $\delta_i > 0$ for all $i \in \{K+1, K+2, ..., n\}$. Then Equation 6 is

$$
\begin{aligned}
&\mathbb{P}(P_{DMT} < u | P_{(n-1)} < u, \boldsymbol{\delta} \succeq 0) \\
=&\left\{ \sum_{i=1}^{K} \left[ u \prod_{j \in \mathcal{K} \setminus i} \frac{u}{2} \right] + (n-K)\left(\frac{u}{2}\right)^K + \psi(n) \left[ \prod_{i=1}^{K} \frac{u}{2} \right] \right\} \Big/ \left\{ \left[ \sum_{i=1}^{n} \prod_{j \in \mathcal{K} \setminus i} u \right] + \psi(n) \prod_{i=1}^{K} u \right\} \\
=&\left\{ uK \left(\frac{u}{2}\right)^{K-1} + (n-K)\left(\frac{u}{2}\right)^K + (1-n)\left(\frac{u}{2}\right)^K \right\} \Big/ \left\{ K u^{K-1} + (n-K)u^K + (1-n)u^K \right\} \\
=&\left(\frac{u}{2}\right)^K (2K + n - K + (1-n)) \Big/ \left\{ u^{K-1}[K + u(n-K) + u(1-n)] \right\} \\
=&\frac{u}{2^K} \left[K+1\right] / \left[K + (1-K)u\right]
\end{aligned}
$$

We can see that this is convex in $u$ because of its similarity to Equation 7 for the origin when we substitute $n$ with $K$ and remove the factors $2^{1-n}$ from Equation 7 and $2^{-K}$ from this expression, which don't affect the derivatives with respect to $u$. So we evaluate the above at its endpoints $u = 0$ where the expression is zero and at $u = \frac{1}{2}$ where it is $\frac{1}{2^K} \leq \frac{1}{2}$ for $K \geq 1$.

**Parameters with mixed signs**

The last scenario under $H_0$ is where $\delta_i > 0$ for at least one $i \in \mathcal{I}$ and $\delta_j < 0$ for at least one $j \in \mathcal{I}$ and $i \neq j$. Then in asymptotic sample sizes for all studies (i.e. $\nu_k \to \infty$ for all $k \in \mathcal{I}$), $\delta_i \to \infty$, $\delta_j \to -\infty$, and

$$\mathbb{P}(P_{DMT} < u | P_{(n-1)} < u) = \mathbb{P}(\hat{S} = 1)\mathbb{P}(P_{(n)} < 2u, | P_{(n-1)} < u) \to 0$$

since $\mathbb{P}(\hat{S} = 1) \leq \mathbb{P}(\hat{\delta}_i > 0, \hat{\delta}_j > 0) + \mathbb{P}(\hat{\delta}_i < 0, \hat{\delta}_j < 0) \to 0$ .

#### Conditional p-value in finite sample sizes

We have the closed form expression of the conditional p-value $\mathbb{P}(P_{DMT} < u | P_{(n-1)} < u)$ in Equation 6 for finite sample sizes. We use this equation to plot the conditional p-value for $n = 3$ in different cases where $\delta_1 = 0$ in the main paper. We see that the conditional probability equals $u$ where $\boldsymbol{\delta} = 0$ or $\delta_3 \neq 0$ and $\nu_3 \to \infty$ for $\delta_2 \geq 0$, which is consistent with our proof when all sample sizes grow unbounded. Whereas, in some situations with finite sample sizes the conditional probability shows very slight inflation, like we found for $n = 2$.
